## Supplementary figures and images for "An efficient behavioral screening platform classifies natural products and other chemical cues according to their chemosensory valence in *C. elegans*"

### Supplemental Figure 1

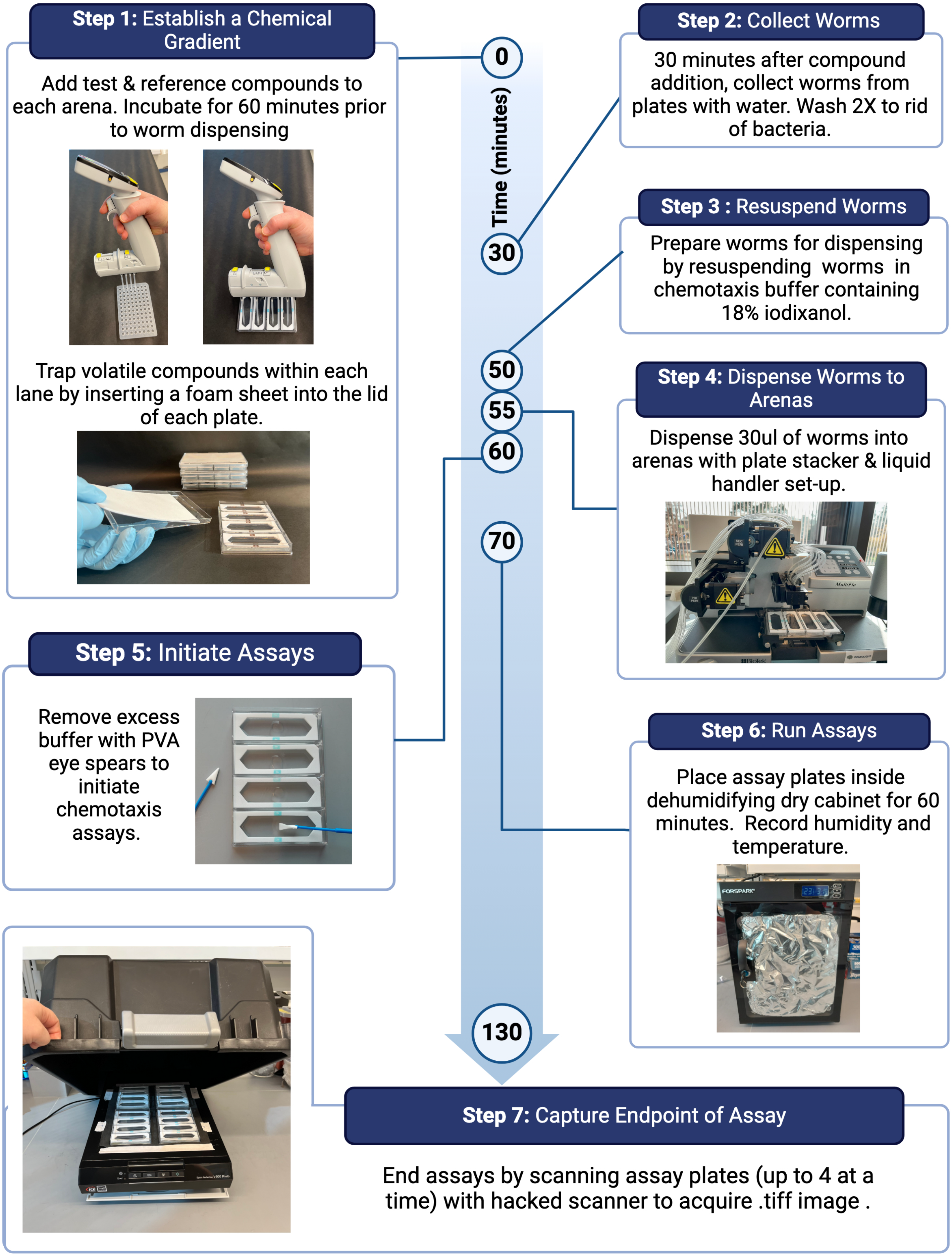

### Supplemental Figure 2

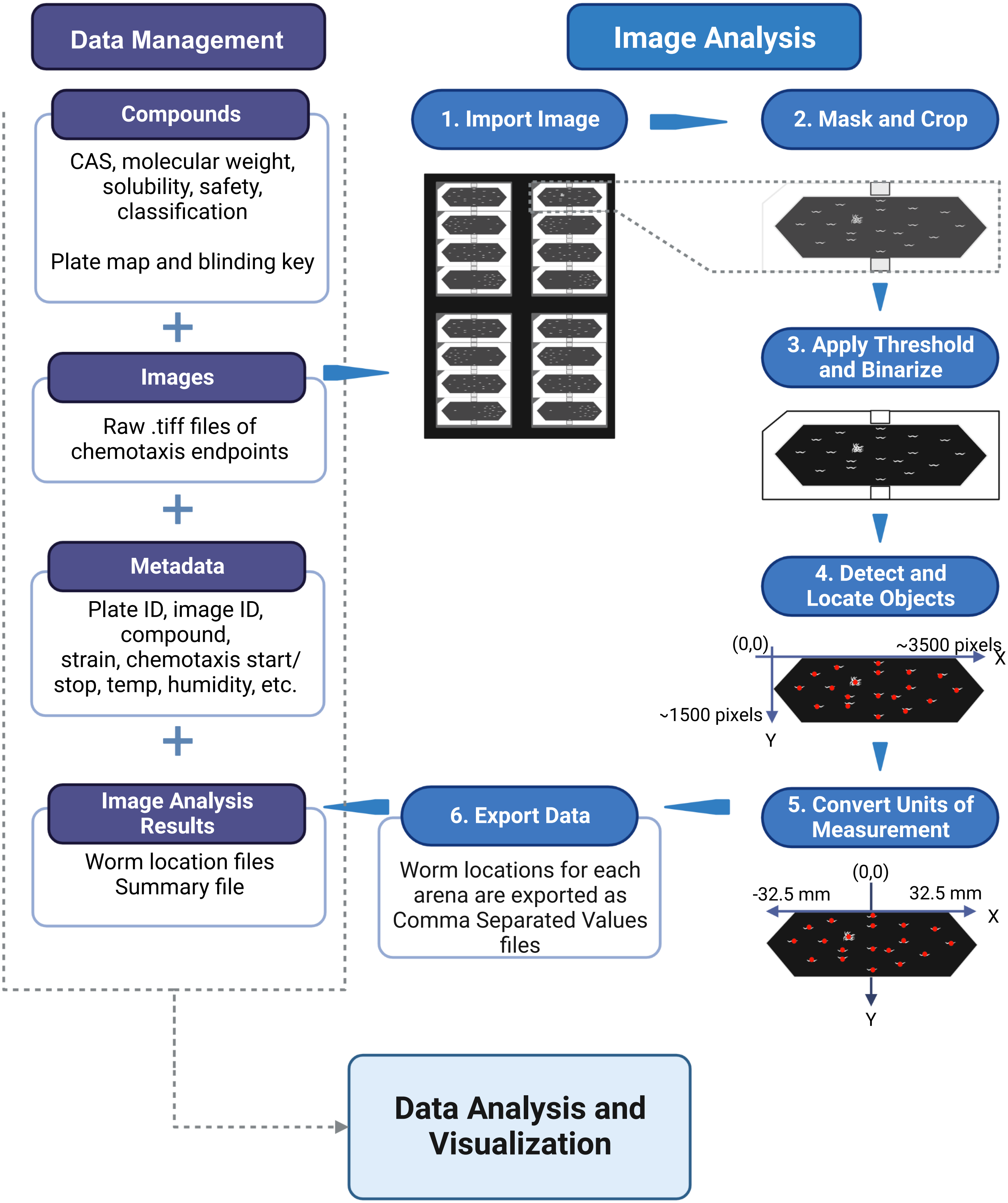
