## Supplemental Table 1 for "An efficient behavioral screening platform classifies natural products and other chemical cues according to their chemosensory valence in *C. elegans*"

S1 Table

| CAS ID | Compound | Vendor | Catalogue # |
|---|---|---|---|
| 0431-03-08 | Diacetyl | TCI | B0682 |
| 0483-04-5 | Ajmalicine | Cayman | 31213 |
| 0491-09-8 | Piperitenone | Cayman | 25752 |
| 100-09-4 | p-Anisic acid | MCE | HY-N1394 |
| 100-66-3 | Anisole | Sigma | 123226 |
| 102518-79-6 | (-)-Huperzine A | MCE | HY-W019711 |
| 104-54-1 | Cinnamyl Alcohol | MCE | HY-Y0078 |
| 104-87-0 | p-Tolualdehyde | MCE | HY-W012860 |
| 105-87-3 | Geranyl Acetate | Ambeed | A404343 |
| 106-22-9 | Citronellol | MCE | HY-W010201 |
| 106-22-9 | β-Citronellol | TCI | C0370 |
| 107-35-7 | Taurine | MCE | HY-B0351 |
| 110-02-1 | Thiophene | TCI | T0223 |
| 111-87-5 | 1-octanol | Sigma | 297877 |
| 112-14-1 | Octyle acetate | TCI | A0042 |
| 112-39-0 | Methyl palmitate | TCI | P0006 |
| 116-26-7 | Safranal | MCE | HY-N7560 |
| 1180-71-8 | Limonin | MCE | HY-17411 |
| 119-65-3 | Isoquinoline | MCE | HY-W012732 |
| 120-57-0 | Piperonyl Alcohol | Sigma | P49104 |
| 123-11-5 | p-anisaldehyde | TCI | A1674 |
| 123-35-3 | Myrcene | Sigma | M100005 |
| 123-51-3 | Isoamyl alcohol | Sigma | W205710 |
| 124-20-9 | Spermidine | MCE | HY-B1776 |
| 126-17-0 | Solasodine (p) | MCE | HY-N0068 |
| 137-32-6 | 2-Methyl-1-butanol | Sigma | 133051 |
| 14371-10-9 | trans-Cinnamaldehyde | MCE | HY-W019711 |
| 1490-04-6 | Menthol | MCE | HY-N1369 |
| 150-86-7 | Phytol | MCE | HY-N3075 |
| 16409-43-1 | L-Mimosine | Apex | B4751 |
| 168316-95-8 | Spinosad (p) | Cayman | 25649 |
| 18524-94-2 | Loganin | MCE | HY-N0512 |
| 18836-52-7 | Pellitorine (p) | Cayman | 11662 |
| 19431-84-6 | Curcumenol | MCE | HY-N2259 |
| 20283-92-5 | Rosmarinic acid | MCE | HY-N0529 |
| 2068-78-2 | Vincristine (sulfate) | MCE | HY-N0488 |
| 2244-16-8 | (+)-Carvone | Sigma | 22070 |
| 23180-57-6 | Paeoniflorin | MCE | HY-N0293 |
| 23800-56-8 | Pogostone | MCE | HY-N1416 |
| 24393-56-4 | Ethyl p-methoxycinnamate | Cayman | 11740 |
| 24697-74-3 | Leonurine | MCE | HY-N0741 |
| 2482-00-0 | Agmatine | Chemimpex | 10668 |
| 3387-41-5 | Sabinene | MCE | HY-108943 |
| 357-70-0 | Galanthamine | MCE | HY-76299 |
| 3650-09-7 | Carnosic acid | MCE | HY-N0644 |
| 372-75-8 | L-Citrulline | MCE | HY-N0391 |
| 37839-63-7 | Germacrene D | Aobious | CFN93281 |
| 4180-23-8 | Trans-Anethole | MCE | HY-N0367 |

| CAS ID | Compound | | Vendor | Catalogue # |
|---|---|---|---|---|
| 4373-41-5 | Maslinic acid | (p) | MCE | HY-N0629 |
| 462-94-2 | Cadaverine | | Sigma | 52063 |
| 464-45-9 | (-)-Borneol | | TCI | B1012 |
| 4674-50-4 | Nootkatone | | MCE | HY-N2195 |
| 469-61-4 | (-)-Cedrene | | MCE | HY-135190 |
| 469-83-0 | Cafestol | | MCE | HY-N6257 |
| 470-82-6 | Eucalyptol | | Sigma | C80601 |
| 474-58-8 | Daucosterol | (p) | MCE | HY-N0410 |
| 474-58-8 | Sitogluside | (p) | Targetmol | T3871 |
| 476-66-4 | Ellagic acid | (p) | MCE | HY-B0183 |
| 484-20-8 | Bergapten | (p) | MCE | HY-N0370 |
| 489-84-9 | Guaiazulene | | Cayman | 31506 |
| 490-79-9 | 2,5-Dihydroxybenzoic acid | | MCE | HY-W001179 |
| 496-16-2 | Coumaran | | MCE | HY-75247 |
| 496-16-2 | 2,3-Dihydrobenzofuran | | TCI | D1583 |
| 499-75-2 | Carvacrol | | MCE | HY-N0711 |
| 508-02-01 | Oleanolic Acid | (p) | MCE | HY-N0156 |
| 520-18-3 | Kaempferol | | MCE | HY-14590 |
| 520-36-5 | Apigenin | (p) | MCE | HY-N1201 |
| 522-17-8 | Deguelin | (p) | MCE | HY-13425 |
| 532-11-6 | Anethole trithione | (p) | MCE | HY-B1223 |
| 536-74-3 | Phenylacetylene | | Sigma | 117706 |
| 5451-09-2 | 5-Aminolevulinic acid (hydrochloride) | | MCE | HY-W000450 |
| 55396-45-7 | 2-Nonylquinolin-4(1H)-one | (p) | Cayman | 9003627 |
| 5784-74-7 | Salsolidine | | MCE | HY-22385 |
| 58-08-2 | Caffeine | | Sigma | C0750 |
| 5957-80-2 | Carnosol | | MCE | HY-N0643 |
| 6080-33-7 | Sinomenine hydrochloride | | MCE | HY-15122 |
| 628-97-7 | Ethyl palmitate | | MCE | HY-N2086 |
| 646-23-1 | Alyssin | | Cayman | 31513 |
| 67-68-5 | DMSO | | VWR | N182 |
| 68370-47-8 | Micheliolide | | MCE | HY-N0847 |
| 689295-71-4 | Salvinorin A Propionate | (p) | Cayman | 22290 |
| 69-72-7 | Salicylic acid | | MCE | HY-B0167 |
| 70-26-8 | L-Ornithine | | MCE | HY-B1352 |
| 7212-44-4 | Nerolidol | | MCE | HY-N1944 |
| 76-22-2 | Camphor | | MCE | HY-N0808 |
| 77-52-1 | Ursolic acid | (p) | MCE | HY-N0140 |
| 821-55-6 | 2-nonanone | | TCI | N093 |
| 83-34-1 | Skatole | | MCE | HY-W007355 |
| 83-79-4 | Rotenone | (p) | MCE | HY-B1756 |
| 84-79-7 | Lapachol | | MCE | HY-N6961 |
| 87-44-5 | Beta caryophyllene | | TCI | C0796 |
| 94-62-2 | Piperine | (p) | MCE | HY-N0144 |
| 98-01-1 | Furfural | | TCI | F0073 |
| 98-86-2 | Acetophenone | | MCE | HY-Y0989 |
| 99-83-2 | α-Phellandrene | | Cayman | 23179 |
| | Water | | NA | NA |
