## Supplemental Table 2 for "An efficient behavioral screening platform classifies natural products and other chemical cues according to their chemosensory valence in *C. elegans*"

| Test_compound | Reference | Test_n | Mean_difference | CI_5 | CI_95 | p_value | BH_correction_0.05 |
|---|---|---|---|---|---|---|---|
| Isoamyl alcohol | DMSO | 502 | 14.167 | 12.82 | 15.512 | 1.35E-94 | 0.0005 |
| 2-Methyl-1-butanol | DMSO | 381 | 11.712 | 9.891 | 13.409 | 6.39E-39 | 0.001 |
| Thiophene | DMSO | 706 | 9.76 | 8.202 | 11.263 | 7.61E-36 | 0.0016 |
| *2,3-Dihydrobenzofuran | DMSO | 705 | 9.194 | 7.63 | 10.674 | 2.43E-32 | 0.0021 |
| Diacetyl | DMSO | 796 | 8.019 | 6.59 | 9.361 | 7.99E-30 | 0.0026 |
| Phenylacetylene | DMSO | 496 | 6.45 | 4.656 | 8.165 | 5.75E-13 | 0.0042 |
| Paeoniflorin | DMSO | 556 | 5.382 | 3.587 | 7.05 | 1.12E-09 | 0.0073 |
| alpha-Phellandrene | DMSO | 777 | 5.368 | 3.801 | 6.836 | 4.13E-12 | 0.0047 |
| Acetophenone | DMSO | 755 | 5.238 | 3.649 | 6.776 | 5.14E-11 | 0.0052 |
| *Coumaran | DMSO | 754 | 4.93 | 3.386 | 6.361 | 8.19E-11 | 0.0057 |
| Leonurine | DMSO | 766 | 4.534 | 3.003 | 6.077 | 7.39E-09 | 0.0078 |
| Guaiazulene | DMSO | 958 | 4.305 | 2.961 | 5.685 | 5.86E-10 | 0.0063 |
| L-Mimosine | DMSO | 916 | 4.282 | 2.806 | 5.731 | 9.49E-09 | 0.0083 |
| Solasodine (p) | DMSO | 826 | 3.821 | 2.257 | 5.414 | 2.10E-06 | 0.0099 |
| Isoquinoline | DMSO | 836 | 3.807 | 2.368 | 5.358 | 5.97E-07 | 0.0094 |
| Furfural | DMSO | 796 | 3.619 | 2.048 | 5.138 | 4.40E-06 | 0.0104 |
| (-)-Huperzine A | DMSO | 911 | 3.378 | 1.87 | 4.772 | 5.08E-06 | 0.0109 |
| Anisole | DMSO | 858 | 3.017 | 1.461 | 4.546 | 1.27E-04 | 0.012 |
| Limonin | DMSO | 907 | 2.823 | 1.348 | 4.346 | 2.23E-04 | 0.0141 |
| Cinnamyl alcohol | DMSO | 857 | 2.814 | 1.287 | 4.297 | 2.47E-04 | 0.0146 |
| Piperitenone | DMSO | 951 | 2.707 | 1.319 | 4.16 | 1.88E-04 | 0.0135 |
| Ethyl palmitate | DMSO | 887 | 2.574 | 1.056 | 4.11 | 9.54E-04 | 0.0161 |
| Carnosol | DMSO | 930 | 2.521 | 1.026 | 3.964 | 7.70E-04 | 0.0151 |
| 2,5-Dihydroxybenzoic acid | DMSO | 971 | 2.463 | 0.976 | 3.874 | 8.65E-04 | 0.0156 |
| p-Tolualdehyde | DMSO | 873 | 2.406 | 0.921 | 3.839 | 0.0012 | 0.0172 |
| Lapachol | DMSO | 811 | 2.354 | 0.861 | 3.925 | 0.0026 | 0.0177 |
| Piperonyl alcohol | DMSO | 1008 | 2.29 | 0.901 | 3.674 | 0.0012 | 0.0167 |
| Sabinene | DMSO | 823 | 2.166 | 0.574 | 3.714 | 0.0069 | 0.0198 |
| Cadaverine | DMSO | 788 | 2.056 | 0.42 | 3.664 | 0.013 | 0.0219 |
| Sinomenine hydrochloride | DMSO | 883 | 2.01 | 0.481 | 3.531 | 0.0098 | 0.0214 |
| (-)-Cedrene | DMSO | 1036 | 2.003 | 0.494 | 3.446 | 0.0078 | 0.0203 |
| Pellitorine (p) | DMSO | 846 | 1.956 | 0.431 | 3.553 | 0.014 | 0.0229 |
| Piperine (p) | DMSO | 787 | 1.949 | 0.41 | 3.518 | 0.014 | 0.0224 |
| ^beta-Citronellol | DMSO | 860 | 1.904 | 0.47 | 3.345 | 0.0094 | 0.0208 |
| Geranyl acetate | DMSO | 890 | 1.807 | 0.275 | 3.329 | 0.0204 | 0.0245 |
| Apigenin (p) | DMSO | 754 | 1.761 | 0.114 | 3.355 | 0.0332 | 0.0266 |
| Salsolidine | DMSO | 748 | 1.706 | 0.113 | 3.244 | 0.0327 | 0.0255 |
| 4-Methoxybenzaldehyde | DMSO | 901 | 1.652 | 0.205 | 3.117 | 0.0262 | 0.025 |
| Nerolidol | DMSO | 781 | 1.556 | 0.112 | 3.041 | 0.0373 | 0.0271 |
| 2-Nonylquinolin-4(1H)-one (p) | DMSO | 803 | 1.511 | 0.051 | 2.933 | 0.0398 | 0.0276 |
| Caffeine | DMSO | 859 | 1.427 | -0.098 | 2.869 | 0.0595 | 0.0281 |
| Agmatine | DMSO | 764 | 1.415 | -0.229 | 2.992 | 0.0852 | 0.0297 |
| Rotenone (p) | DMSO | 865 | 1.405 | -0.153 | 2.948 | 0.0757 | 0.0286 |
| Menthol | DMSO | 928 | 1.359 | -0.159 | 2.899 | 0.0815 | 0.0292 |
| ^Citronellol | DMSO | 767 | 1.28 | -0.217 | 2.766 | 0.0927 | 0.0302 |
| (+)-Carvone | DMSO | 1052 | 1.122 | -0.31 | 2.58 | 0.1279 | 0.0313 |
| Spermidine | DMSO | 977 | 0.904 | -0.559 | 2.389 | 0.2292 | 0.0318 |
| †Sitogluside (p) | DMSO | 993 | 0.802 | -0.787 | 2.277 | 0.3046 | 0.0349 |

| | | | | | | | |
|---|---|---|---|---|---|---|---|
| Ajmalicine | DMSO | 864 | 0.798 | -0.724 | 2.329 | 0.3057 | 0.0354 |
| Maslinic acid (p) | DMSO | 879 | 0.791 | -0.692 | 2.348 | 0.3075 | 0.0365 |
| 5-Aminolevulinic acid (hydrochloride) | DMSO | 838 | 0.779 | -0.658 | 2.288 | 0.3001 | 0.0344 |
| Nootkatone | DMSO | 963 | 0.671 | -0.795 | 2.107 | 0.3645 | 0.0375 |
| Alyssin | DMSO | 970 | 0.601 | -0.857 | 2.071 | 0.421 | 0.0396 |
| Eucalyptol | DMSO | 853 | 0.549 | -1.033 | 2.06 | 0.4862 | 0.0406 |
| Carnosic acid | DMSO | 797 | 0.534 | -1.109 | 2.127 | 0.5174 | 0.0411 |
| Skatole | DMSO | 978 | 0.436 | -1.005 | 1.948 | 0.5629 | 0.0417 |
| Water | DMSO | 915 | 0.418 | -1.043 | 1.889 | 0.5758 | 0.0422 |
| trans-Anethole | DMSO | 963 | 0.346 | -1.117 | 1.851 | 0.6473 | 0.0432 |
| Rosmarinic acid | DMSO | 802 | 0.318 | -1.193 | 1.89 | 0.6859 | 0.0448 |
| Vincristine (sulfate) | DMSO | 625 | 0.23 | -1.497 | 1.953 | 0.7935 | 0.0458 |
| Galanthamine | DMSO | 886 | 0.203 | -1.288 | 1.744 | 0.7931 | 0.0453 |
| L-Citrulline | DMSO | 883 | 0.129 | -1.363 | 1.701 | 0.8688 | 0.0469 |
| Salicylic acid | DMSO | 1058 | 0.082 | -1.329 | 1.516 | 0.9101 | 0.0474 |
| Bergapten (p) | DMSO | 806 | 0.079 | -1.512 | 1.704 | 0.9231 | 0.0479 |
| Deguelin (p) | DMSO | 972 | 0.051 | -1.425 | 1.597 | 0.9472 | 0.049 |
| trans-Cinnamaldehyde | DMSO | 879 | 0.043 | -1.444 | 1.598 | 0.9555 | 0.0495 |
| Curcumenol | DMSO | 1045 | -0.055 | -1.488 | 1.424 | 0.9415 | 0.0484 |
| Taurine | DMSO | 985 | -0.151 | -1.61 | 1.341 | 0.841 | 0.0464 |
| Cafestol | DMSO | 1046 | -0.31 | -1.719 | 1.135 | 0.6699 | 0.0443 |
| Pogostone | DMSO | 856 | -0.356 | -1.886 | 1.211 | 0.6522 | 0.0438 |
| Loganin | DMSO | 928 | -0.403 | -1.956 | 1.024 | 0.5962 | 0.0427 |
| Kaempferol | DMSO | 999 | -0.579 | -2.004 | 0.854 | 0.4269 | 0.0401 |
| L-Ornithine | DMSO | 908 | -0.61 | -2.117 | 0.831 | 0.4174 | 0.0391 |
| Anethole trithione (p) | DMSO | 926 | -0.628 | -2.084 | 0.863 | 0.4035 | 0.0385 |
| Carvacrol | DMSO | 935 | -0.65 | -2.091 | 0.8 | 0.3781 | 0.038 |
| Beta caryophyllene | DMSO | 1041 | -0.75 | -2.169 | 0.597 | 0.2881 | 0.0333 |
| Germacrene D | DMSO | 1158 | -0.754 | -2.252 | 0.595 | 0.2992 | 0.0339 |
| Micheliolide | DMSO | 808 | -0.762 | -2.259 | 0.733 | 0.3184 | 0.037 |
| (-)-Borneol | DMSO | 867 | -0.793 | -2.327 | 0.716 | 0.3073 | 0.0359 |
| Octyle acetate | DMSO | 859 | -0.875 | -2.397 | 0.677 | 0.2646 | 0.0328 |
| Myrcene | DMSO | 769 | -0.923 | -2.505 | 0.62 | 0.2472 | 0.0323 |
| p-Anisic acid | DMSO | 1024 | -1.168 | -2.624 | 0.314 | 0.1191 | 0.0307 |
| Oleanolic acid (p) | DMSO | 730 | -1.76 | -3.362 | -0.125 | 0.033 | 0.026 |
| †Daucosterol (p) | DMSO | 804 | -1.885 | -3.467 | -0.321 | 0.0189 | 0.024 |
| Ethyl p-methoxycinnamate | DMSO | 751 | -1.895 | -3.396 | -0.335 | 0.0152 | 0.0234 |
| Methyl palmitate | DMSO | 935 | -1.978 | -3.418 | -0.567 | 0.0065 | 0.0193 |
| Safranal | DMSO | 848 | -2.081 | -3.542 | -0.589 | 0.0057 | 0.0188 |
| Ursolic acid (p) | DMSO | 884 | -2.229 | -3.737 | -0.737 | 0.0036 | 0.0182 |
| Camphor | DMSO | 941 | -2.756 | -4.166 | -1.342 | 1.30E-04 | 0.0125 |
| Spinosad (p) | DMSO | 900 | -2.797 | -4.28 | -1.379 | 1.57E-04 | 0.013 |
| Salvinorin A propionate (p) | DMSO | 926 | -3.313 | -4.793 | -1.847 | 1.04E-05 | 0.0115 |
| Ellagic acid (p) | DMSO | 890 | -3.744 | -5.184 | -2.379 | 1.68E-07 | 0.0089 |
| 2-Nonanone | DMSO | 867 | -4.576 | -6.02 | -3.097 | 8.42E-10 | 0.0068 |
| Phytol | DMSO | 625 | -6.249 | -7.8 | -4.623 | 1.27E-14 | 0.0036 |
| 1-Octanol | DMSO | 652 | -7.446 | -8.945 | -5.917 | 5.39E-22 | 0.0031 |
| Isoamyl alcohol | H2O | 502 | 13.749 | 12.342 | 15.152 | 5.67E-82 | 0.0005 |
| 2-Methyl-1-butanol | H2O | 381 | 11.294 | 9.483 | 13.039 | 1.41E-35 | 0.001 |

| | | | | | | | |
|---|---|---|---|---|---|---|---|
| Thiophene | H2O | 706 | 9.341 | 7.723 | 10.903 | 1.12E-30 | 0.0016 |
| *2,3-Dihydrobenzofuran | H2O | 705 | 8.776 | 7.138 | 10.356 | 1.12E-26 | 0.0021 |
| Diacetyl | H2O | 796 | 7.6 | 6.171 | 9.055 | 5.01E-25 | 0.0026 |
| Phenylacetylene | H2O | 496 | 6.032 | 4.136 | 7.734 | 4.99E-11 | 0.0042 |
| Paeoniflorin | H2O | 556 | 4.963 | 3.159 | 6.769 | 7.07E-08 | 0.0073 |
| alpha-Phellandrene | H2O | 777 | 4.949 | 3.271 | 6.52 | 2.34E-09 | 0.0052 |
| Acetophenone | H2O | 755 | 4.82 | 3.18 | 6.423 | 5.69E-09 | 0.0057 |
| *Coumaran | H2O | 754 | 4.512 | 2.942 | 6.024 | 9.54E-09 | 0.0063 |
| Leonurine | H2O | 766 | 4.115 | 2.436 | 5.682 | 6.67E-07 | 0.0089 |
| Guaiazulene | H2O | 958 | 3.886 | 2.422 | 5.379 | 2.60E-07 | 0.0078 |
| L-Mimosine | H2O | 916 | 3.864 | 2.354 | 5.385 | 5.83E-07 | 0.0083 |
| Solasodine (p) | H2O | 826 | 3.402 | 1.843 | 4.984 | 2.17E-05 | 0.0099 |
| Isoquinoline | H2O | 836 | 3.389 | 1.76 | 4.888 | 2.17E-05 | 0.0104 |
| Furfural | H2O | 796 | 3.201 | 1.56 | 4.793 | 1.04E-04 | 0.012 |
| (-)-Huperzine A | H2O | 911 | 2.96 | 1.392 | 4.435 | 1.38E-04 | 0.0125 |
| Anisole | H2O | 858 | 2.598 | 0.981 | 4.142 | 0.0013 | 0.0141 |
| Limonin | H2O | 907 | 2.404 | 0.847 | 3.964 | 0.0025 | 0.0151 |
| Cinnamyl alcohol | H2O | 857 | 2.395 | 0.771 | 3.941 | 0.0031 | 0.0156 |
| Piperitenone | H2O | 951 | 2.288 | 0.791 | 3.827 | 0.0031 | 0.0161 |
| Ethyl palmitate | H2O | 887 | 2.156 | 0.531 | 3.782 | 0.0093 | 0.0193 |
| Carnosol | H2O | 930 | 2.102 | 0.485 | 3.611 | 0.0084 | 0.0182 |
| 2,5-Dihydroxybenzoic acid | H2O | 971 | 2.045 | 0.514 | 3.517 | 0.0076 | 0.0177 |
| p-Tolualdehyde | H2O | 873 | 1.987 | 0.397 | 3.483 | 0.0116 | 0.0203 |
| Lapachol | H2O | 811 | 1.936 | 0.308 | 3.582 | 0.0205 | 0.0208 |
| Piperonyl alcohol | H2O | 1008 | 1.872 | 0.43 | 3.297 | 0.0105 | 0.0198 |
| Sabinene | H2O | 823 | 1.747 | 0.129 | 3.39 | 0.0357 | 0.0214 |
| Cadaverine | H2O | 788 | 1.637 | -0.068 | 3.252 | 0.0532 | 0.0234 |
| Sinomenine hydrochloride | H2O | 883 | 1.592 | 0.016 | 3.156 | 0.0469 | 0.0229 |
| (-)-Cedrene | H2O | 1036 | 1.584 | -0.021 | 3.08 | 0.0452 | 0.0224 |
| Pellitorine (p) | H2O | 846 | 1.538 | -0.211 | 3.091 | 0.0679 | 0.025 |
| Piperine (p) | H2O | 787 | 1.531 | -0.104 | 3.112 | 0.0621 | 0.0245 |
| ^beta-Citronellol | H2O | 860 | 1.486 | -0.115 | 2.991 | 0.0608 | 0.024 |
| Geranyl acetate | H2O | 890 | 1.388 | -0.257 | 2.948 | 0.0895 | 0.0255 |
| Apigenin (p) | H2O | 754 | 1.342 | -0.304 | 3.018 | 0.1132 | 0.0271 |
| Salsolidine | H2O | 748 | 1.287 | -0.364 | 2.899 | 0.122 | 0.0292 |
| 4-Methoxybenzaldehyde | H2O | 901 | 1.233 | -0.367 | 2.726 | 0.118 | 0.0281 |
| Nerolidol | H2O | 781 | 1.138 | -0.413 | 2.689 | 0.1505 | 0.0307 |
| 2-Nonylquinolin-4(1H)-one (p) | H2O | 803 | 1.093 | -0.502 | 2.622 | 0.1704 | 0.0313 |
| Caffeine | H2O | 859 | 1.008 | -0.528 | 2.521 | 0.1949 | 0.0339 |
| Agmatine | H2O | 764 | 0.996 | -0.63 | 2.591 | 0.2254 | 0.0349 |
| Rotenone (p) | H2O | 865 | 0.986 | -0.654 | 2.512 | 0.2219 | 0.0344 |
| Menthol | H2O | 928 | 0.941 | -0.662 | 2.471 | 0.2392 | 0.0354 |
| ^Citronellol | H2O | 767 | 0.861 | -0.705 | 2.391 | 0.2755 | 0.0359 |
| (+)-Carvone | H2O | 1052 | 0.704 | -0.733 | 2.151 | 0.3388 | 0.0375 |
| Spermidine | H2O | 977 | 0.486 | -1.046 | 2.016 | 0.5339 | 0.0391 |
| †Sitogluside (p) | H2O | 993 | 0.384 | -1.175 | 1.947 | 0.6297 | 0.0406 |
| Ajmalicine | H2O | 864 | 0.379 | -1.246 | 1.947 | 0.6417 | 0.0417 |
| Maslinic acid (p) | H2O | 879 | 0.373 | -1.208 | 1.963 | 0.6448 | 0.0427 |
| 5-Aminolevulinic acid (hydrochloride) | H2O | 838 | 0.36 | -1.164 | 1.883 | 0.6431 | 0.0422 |

| | | | | | | | |
|---|---|---|---|---|---|---|---|
| Nootkatone | H2O | 963 | 0.253 | -1.31 | 1.695 | 0.7414 | 0.0453 |
| Alyssin | H2O | 970 | 0.183 | -1.355 | 1.693 | 0.8141 | 0.0464 |
| Eucalyptol | H2O | 853 | 0.131 | -1.52 | 1.711 | 0.8737 | 0.0474 |
| Carnosic acid | H2O | 797 | 0.116 | -1.503 | 1.752 | 0.8888 | 0.0479 |
| Skatole | H2O | 978 | 0.017 | -1.501 | 1.572 | 0.9822 | 0.0495 |
| trans-Anethole | H2O | 963 | -0.072 | -1.68 | 1.456 | 0.9282 | 0.049 |
| Rosmarinic acid | H2O | 802 | -0.1 | -1.654 | 1.485 | 0.9003 | 0.0484 |
| Vincristine (sulfate) | H2O | 625 | -0.188 | -1.963 | 1.556 | 0.8341 | 0.0469 |
| Galanthamine | H2O | 886 | -0.216 | -1.769 | 1.336 | 0.7856 | 0.0458 |
| L-Citrulline | H2O | 883 | -0.289 | -1.882 | 1.252 | 0.7175 | 0.0448 |
| Salicylic acid | H2O | 1058 | -0.336 | -1.861 | 1.087 | 0.6546 | 0.0438 |
| Bergapten (p) | H2O | 806 | -0.339 | -2.005 | 1.31 | 0.6883 | 0.0443 |
| Deguelin (p) | H2O | 972 | -0.367 | -1.87 | 1.199 | 0.639 | 0.0411 |
| trans-Cinnamaldehyde | H2O | 879 | -0.375 | -1.989 | 1.223 | 0.6472 | 0.0432 |
| Curcumenol | H2O | 1068 | -0.473 | -1.981 | 1.022 | 0.537 | 0.0396 |
| Taurine | H2O | 1045 | -0.569 | -2.095 | 1.017 | 0.4732 | 0.0385 |
| Cafestol | H2O | 985 | -0.729 | -2.238 | 0.679 | 0.3274 | 0.037 |
| Pogostone | H2O | 1046 | -0.775 | -2.391 | 0.809 | 0.3428 | 0.038 |
| Loganin | H2O | 856 | -0.821 | -2.41 | 0.684 | 0.2982 | 0.0365 |
| Kaempferol | H2O | 928 | -0.998 | -2.493 | 0.486 | 0.1894 | 0.0328 |
| L-Ornithine | H2O | 999 | -1.028 | -2.636 | 0.465 | 0.1937 | 0.0333 |
| Anethole trithione (p) | H2O | 908 | -1.047 | -2.616 | 0.412 | 0.1755 | 0.0323 |
| Carvacrol | H2O | 926 | -1.068 | -2.638 | 0.43 | 0.1723 | 0.0318 |
| Beta caryophyllene | H2O | 935 | -1.168 | -2.683 | 0.244 | 0.1178 | 0.0276 |
| Germacrene D | H2O | 1041 | -1.172 | -2.614 | 0.272 | 0.1113 | 0.0266 |
| Micheliolide | H2O | 1158 | -1.18 | -2.724 | 0.364 | 0.1341 | 0.0302 |
| (-)-Borneol | H2O | 808 | -1.211 | -2.782 | 0.362 | 0.1311 | 0.0297 |
| Octyle acetate | H2O | 867 | -1.293 | -2.994 | 0.255 | 0.1186 | 0.0286 |
| Myrcene | H2O | 859 | -1.341 | -2.95 | 0.272 | 0.1028 | 0.026 |
| p-Anisic acid | H2O | 769 | -1.587 | -3.143 | -0.052 | 0.0442 | 0.0219 |
| Oleanolic acid (p) | H2O | 1024 | -2.179 | -3.833 | -0.584 | 0.0086 | 0.0188 |
| †Daucosterol (p) | H2O | 730 | -2.303 | -3.945 | -0.635 | 0.0064 | 0.0172 |
| Ethyl p-methoxycinnamate | H2O | 804 | -2.314 | -3.919 | -0.7 | 0.0049 | 0.0167 |
| Methyl palmitate | H2O | 751 | -2.397 | -3.896 | -0.89 | 0.0018 | 0.0146 |
| Safranal | H2O | 935 | -2.5 | -4.01 | -1.003 | 0.0011 | 0.0135 |
| Ursolic acid (p) | H2O | 848 | -2.648 | -4.195 | -1.07 | 8.98E-04 | 0.013 |
| Camphor | H2O | 884 | -3.175 | -4.717 | -1.696 | 3.81E-05 | 0.0115 |
| Spinosad (p) | H2O | 941 | -3.215 | -4.792 | -1.771 | 3.01E-05 | 0.0109 |
| Salvinorin A propionate (p) | H2O | 900 | -3.732 | -5.301 | -2.233 | 1.86E-06 | 0.0094 |
| Ellagic acid (p) | H2O | 926 | -4.162 | -5.656 | -2.786 | 1.31E-08 | 0.0068 |
| 2-Nonanone | H2O | 890 | -4.994 | -6.571 | -3.473 | 2.63E-10 | 0.0047 |
| Phytol | H2O | 867 | -6.667 | -8.326 | -4.999 | 4.00E-15 | 0.0036 |
| 1-Octanol | H2O | 625 | -7.865 | -9.412 | -6.308 | 3.08E-23 | 0.0031 |
