## Supplemental Table 3 for "An efficient behavioral screening platform classifies natural products and other chemical cues according to their chemosensory valence in *C. elegans*"

| Test compound | Control | Test_n | Mean difference | CI_5 | CI_95 | Strain |
|---|---|---|---|---|---|---|
| (-)-Huperzine A | DMSO | 675 | 2.507 | 1.196 | 3.777 | PR678 |
| 1-octanol | DMSO | 668 | -3.953 | -5.293 | -2.707 | PR678 |
| 2,3-Dihydrobenzofuran | DMSO | 670 | 6.791 | 5.382 | 8.233 | PR678 |
| 2,5-Dihydroxybenzoic acid | DMSO | 998 | 1.257 | 0.015 | 2.37 | PR678 |
| 2-Methyl-1-butanol | DMSO | 640 | 1.723 | 0.325 | 3.059 | PR678 |
| 2-nonanone | DMSO | 767 | -6.326 | -7.514 | -5.134 | PR678 |
| Acetophenone | DMSO | 636 | -4.155 | -5.44 | -2.91 | PR678 |
| Anisole | DMSO | 777 | 6.443 | 5.207 | 7.7 | PR678 |
| Camphor | DMSO | 624 | 1.511 | 0.138 | 2.831 | PR678 |
| Carnosol | DMSO | 635 | 6.716 | 5.223 | 8.15 | PR678 |
| Cinnamyl alcohol | DMSO | 727 | 4.824 | 3.422 | 6.117 | PR678 |
| Coumaran | DMSO | 808 | 4.437 | 3.143 | 5.693 | PR678 |
| Daucosterol | DMSO | 673 | 4.354 | 3.02 | 5.66 | PR678 |
| Diacetyl | DMSO | 738 | 12.024 | 10.764 | 13.222 | PR678 |
| Ellagic acid | DMSO | 671 | 3.74 | 2.417 | 5.034 | PR678 |
| Ethyl p-methoxycinnamate | DMSO | 620 | 0.164 | -1.115 | 1.435 | PR678 |
| Ethyl palmitate | DMSO | 681 | 2.073 | 0.764 | 3.35 | PR678 |
| Furfural | DMSO | 574 | -1.76 | -3.071 | -0.422 | PR678 |
| Guaiazulene | DMSO | 593 | 1.399 | 0.053 | 2.774 | PR678 |
| Water | DMSO | 523 | 3.816 | 2.391 | 5.156 | PR678 |
| Isoamyl alcohol | DMSO | 889 | 3.361 | 2.089 | 4.599 | PR678 |
| Isoquinoline | DMSO | 582 | 1.29 | 0.012 | 2.51 | PR678 |
| L-Mimosine | DMSO | 884 | 1.955 | 0.725 | 3.121 | PR678 |
| Lapachol | DMSO | 640 | 3.359 | 2.046 | 4.713 | PR678 |
| Leonurine | DMSO | 555 | 3.117 | 1.725 | 4.466 | PR678 |
| Limonin | DMSO | 782 | 2.044 | 0.741 | 3.283 | PR678 |
| Methyl palmitate | DMSO | 798 | 3.058 | 1.888 | 4.282 | PR678 |
| Oleanolic acid | DMSO | 617 | 5.09 | 3.591 | 6.546 | PR678 |
| Paeoniflorin | DMSO | 595 | 2.592 | 1.236 | 3.957 | PR678 |
| Phenylacetylene | DMSO | 794 | 3.332 | 2.119 | 4.579 | PR678 |
| Phytol | DMSO | 626 | 4.162 | 2.887 | 5.405 | PR678 |
| Piperitenone | DMSO | 863 | 5.808 | 4.469 | 7.081 | PR678 |
| Piperonyl Alcohol | DMSO | 615 | 2.855 | 1.46 | 4.164 | PR678 |
| Sabinene | DMSO | 816 | 3.51 | 2.242 | 4.766 | PR678 |
| Salvinorin A Propionate | DMSO | 714 | 2.695 | 1.401 | 3.977 | PR678 |
| Sinomenine hydrochloride | DMSO | 673 | 3.462 | 2.183 | 4.745 | PR678 |
| Solasodine | DMSO | 790 | 4.739 | 3.416 | 5.97 | PR678 |
| Spinosad | DMSO | 846 | 2.191 | 0.973 | 3.418 | PR678 |
| Thiophene | DMSO | 620 | 2.925 | 1.572 | 4.242 | PR678 |
| Ursolic acid | DMSO | 559 | 3.597 | 2.163 | 4.994 | PR678 |
| p-Tolualdehyde | DMSO | 734 | -0.583 | -1.815 | 0.656 | PR678 |
| alpha-Phellandrene | DMSO | 879 | 2.643 | 1.416 | 3.875 | PR678 |
| (-)-Huperzine A | DMSO | 976 | 0.283 | -1.202 | 1.854 | CX10 |

| | | | | | | |
|---|---|---|---|---|---|---|
| 1-octanol | DMSO | 806 | -1.173 | -2.655 | 0.347 | CX10 |
| 2,3-Dihydrobenzofuran | DMSO | 796 | 9.736 | 8.303 | 11.154 | CX10 |
| 2,5-Dihydroxybenzoic acid | DMSO | 840 | -1.52 | -3 | 0.088 | CX10 |
| 2-Methyl-1-butanol | DMSO | 558 | 10.784 | 9.256 | 12.363 | CX10 |
| 2-nonanone | DMSO | 861 | -3.745 | -5.229 | -2.16 | CX10 |
| Acetophenone | DMSO | 1056 | 1.139 | -0.365 | 2.594 | CX10 |
| Anisole | DMSO | 760 | 4.987 | 3.459 | 6.677 | CX10 |
| Camphor | DMSO | 1049 | 1.568 | 0.083 | 3.055 | CX10 |
| Carnosol | DMSO | 974 | 1.664 | 0.193 | 3.204 | CX10 |
| Cinnamyl alcohol | DMSO | 1007 | 1.396 | -0.085 | 2.902 | CX10 |
| Coumaran | DMSO | 987 | 2.801 | 1.304 | 4.346 | CX10 |
| Daucosterol | DMSO | 1002 | 0.196 | -1.364 | 1.754 | CX10 |
| Diacetyl | DMSO | 861 | 3.998 | 2.468 | 5.489 | CX10 |
| Ellagic acid | DMSO | 1042 | 0.994 | -0.526 | 2.439 | CX10 |
| Ethyl p-methoxycinnamate | DMSO | 894 | 0.882 | -0.538 | 2.276 | CX10 |
| Ethyl palmitate | DMSO | 815 | 0.388 | -1.19 | 1.952 | CX10 |
| Furfural | DMSO | 780 | 3.522 | 1.92 | 5.09 | CX10 |
| Guaiazulene | DMSO | 861 | -2.027 | -3.497 | -0.525 | CX10 |
| Water | DMSO | 909 | 2.071 | 0.554 | 3.541 | CX10 |
| Isoamyl alcohol | DMSO | 711 | 11.387 | 9.916 | 12.882 | CX10 |
| Isoquinoline | DMSO | 994 | 1.71 | 0.223 | 3.175 | CX10 |
| L-Mimosine | DMSO | 855 | 0.684 | -0.812 | 2.197 | CX10 |
| Lapachol | DMSO | 1126 | 0.36 | -1.049 | 1.817 | CX10 |
| Leonurine | DMSO | 1061 | -0.263 | -1.726 | 1.199 | CX10 |
| Limonin | DMSO | 973 | 2.249 | 0.792 | 3.709 | CX10 |
| Methyl palmitate | DMSO | 952 | 1.587 | 0.087 | 3.159 | CX10 |
| Oleanolic acid | DMSO | 908 | -1.868 | -3.371 | -0.368 | CX10 |
| Paeoniflorin | DMSO | 825 | 4.498 | 2.976 | 6.031 | CX10 |
| Phenylacetylene | DMSO | 768 | 5.656 | 4.108 | 7.252 | CX10 |
| Phytol | DMSO | 755 | 0.505 | -1.089 | 2.096 | CX10 |
| Piperitenone | DMSO | 875 | 4.663 | 3.171 | 6.127 | CX10 |
| Piperonyl Alcohol | DMSO | 962 | -4.272 | -5.744 | -2.766 | CX10 |
| Sabinene | DMSO | 939 | 1.9 | 0.345 | 3.413 | CX10 |
| Salvinorin A Propionate | DMSO | 964 | -0.629 | -2.084 | 1 | CX10 |
| Sinomenine hydrochloride | DMSO | 1028 | -1.037 | -2.516 | 0.471 | CX10 |
| Solasodine | DMSO | 1051 | -0.543 | -2.039 | 0.908 | CX10 |
| Spinosad | DMSO | 895 | 0.993 | -0.553 | 2.536 | CX10 |
| Thiophene | DMSO | 897 | 1.565 | -0.022 | 3.119 | CX10 |
| Ursolic acid | DMSO | 869 | 0.112 | -1.384 | 1.615 | CX10 |
| p-Tolualdehyde | DMSO | 954 | 2.369 | 0.88 | 3.945 | CX10 |
| alpha-Phellandrene | DMSO | 962 | 2.3 | 0.835 | 3.801 | CX10 |
| (-)-Huperzine A | DMSO | 596 | -0.435 | -1.591 | 0.675 | GN1077 |
| 1-octanol | DMSO | 811 | -0.041 | -0.936 | 0.853 | GN1077 |
| 2,3-Dihydrobenzofuran | DMSO | 851 | -1.425 | -2.399 | -0.482 | GN1077 |

| | | | | | | |
|---|---|---|---|---|---|---|
| 2,5-Dihydroxybenzoic acid | DMSO | 1024 | -0.758 | -1.629 | 0.081 | GN1077 |
| 2-Methyl-1-butanol | DMSO | 434 | -2.09 | -3.444 | -0.752 | GN1077 |
| 2-nonanone | DMSO | 709 | -2.109 | -3.044 | -1.217 | GN1077 |
| Acetophenone | DMSO | 830 | -2.663 | -3.556 | -1.767 | GN1077 |
| Anisole | DMSO | 668 | -0.644 | -1.665 | 0.35 | GN1077 |
| Camphor | DMSO | 857 | -0.214 | -1.216 | 0.732 | GN1077 |
| Carnosol | DMSO | 693 | -0.379 | -1.344 | 0.546 | GN1077 |
| Cinnamyl alcohol | DMSO | 554 | -0.529 | -1.719 | 0.685 | GN1077 |
| Coumaran | DMSO | 841 | -1.106 | -2.007 | -0.226 | GN1077 |
| Daucosterol | DMSO | 642 | 1.867 | 0.797 | 2.996 | GN1077 |
| Diacetyl | DMSO | 836 | -0.553 | -1.417 | 0.351 | GN1077 |
| Ellagic acid | DMSO | 887 | -0.999 | -1.885 | -0.117 | GN1077 |
| Ethyl p-methoxycinnamate | DMSO | 784 | 0.348 | -0.571 | 1.248 | GN1077 |
| Ethyl palmitate | DMSO | 961 | -1.321 | -2.299 | -0.361 | GN1077 |
| Furfural | DMSO | 733 | -1.74 | -2.628 | -0.835 | GN1077 |
| Guaiazulene | DMSO | 873 | -0.761 | -1.726 | 0.256 | GN1077 |
| Water | DMSO | 784 | -1.167 | -2.06 | -0.277 | GN1077 |
| Isoamyl alcohol | DMSO | 703 | -1.151 | -2.043 | -0.251 | GN1077 |
| Isoquinoline | DMSO | 732 | -1.789 | -2.652 | -0.943 | GN1077 |
| L-Mimosine | DMSO | 700 | 0.141 | -0.777 | 1.026 | GN1077 |
| Lapachol | DMSO | 632 | -1.912 | -2.954 | -0.914 | GN1077 |
| Leonurine | DMSO | 619 | -0.248 | -1.239 | 0.691 | GN1077 |
| Limonin | DMSO | 722 | -0.283 | -1.295 | 0.713 | GN1077 |
| Methyl palmitate | DMSO | 656 | -1.244 | -2.16 | -0.294 | GN1077 |
| Oleanolic acid | DMSO | 675 | -0.558 | -1.557 | 0.442 | GN1077 |
| Paeoniflorin | DMSO | 664 | -1.207 | -2.325 | -0.14 | GN1077 |
| Phenylacetylene | DMSO | 495 | -1.36 | -2.34 | -0.331 | GN1077 |
| Phytol | DMSO | 678 | -1.793 | -2.723 | -0.88 | GN1077 |
| Piperitenone | DMSO | 747 | -0.641 | -1.638 | 0.328 | GN1077 |
| Piperonyl Alcohol | DMSO | 745 | -1.812 | -2.737 | -0.91 | GN1077 |
| Sabinene | DMSO | 681 | -1.018 | -2.02 | 0.006 | GN1077 |
| Salvinorin A Propionate | DMSO | 714 | -1.218 | -2.163 | -0.303 | GN1077 |
| Sinomenine hydrochloride | DMSO | 897 | -1.688 | -2.576 | -0.775 | GN1077 |
| Solasodine | DMSO | 702 | 1.259 | 0.214 | 2.336 | GN1077 |
| Spinosad | DMSO | 764 | 0.921 | -0.017 | 1.928 | GN1077 |
| Thiophene | DMSO | 861 | -1.322 | -2.201 | -0.448 | GN1077 |
| Ursolic acid | DMSO | 868 | -1.521 | -2.45 | -0.639 | GN1077 |
| p-Tolualdehyde | DMSO | 824 | -0.262 | -1.151 | 0.663 | GN1077 |
| alpha-Phellandrene | DMSO | 764 | -1.078 | -2.012 | -0.176 | GN1077 |
