## Supplemental Table 4 for "An efficient behavioral screening platform classifies natural products and other chemical cues according to their chemosensory valence in *C. elegans*"

| Test compound | CI_5 | CI_95 | Delta-delta | Strain 1 | Strain 2 |
|---|---|---|---|---|---|
| (-)-Huperzine A | -5.646 | -1.986 | -3.813 | GN1077 | N2 |
| 1-octanol | 5.703 | 9.104 | 7.406 | GN1077 | N2 |
| 2,3-Dihydrobenzofuran | -12.331 | -8.728 | -10.619 | GN1077 | N2 |
| 2,5-Dihydroxybenzoic acid | -4.863 | -1.469 | -3.221 | GN1077 | N2 |
| 2-Methyl-1-butanol | -15.918 | -11.621 | -13.802 | GN1077 | N2 |
| 2-nonanone | 0.650 | 4.167 | 2.467 | GN1077 | N2 |
| Acetophenone | -9.646 | -6.084 | -7.901 | GN1077 | N2 |
| Anisole | -5.446 | -1.800 | -3.661 | GN1077 | N2 |
| Camphor | 0.892 | 4.240 | 2.542 | GN1077 | N2 |
| Carnosol | -4.629 | -1.139 | -2.899 | GN1077 | N2 |
| Cinnamyl alcohol | -5.274 | -1.386 | -3.342 | GN1077 | N2 |
| Coumaran | -7.705 | -4.306 | -6.037 | GN1077 | N2 |
| Daucosterol | 1.864 | 5.701 | 3.751 | GN1077 | N2 |
| Diacetyl | -10.197 | -6.903 | -8.572 | GN1077 | N2 |
| Ellagic acid | 1.123 | 4.446 | 2.745 | GN1077 | N2 |
| Ethyl p-methoxycinnamate | 0.497 | 4.095 | 2.243 | GN1077 | N2 |
| Ethyl palmitate | -5.656 | -2.052 | -3.895 | GN1077 | N2 |
| Furfural | -7.098 | -3.556 | -5.359 | GN1077 | N2 |
| Guaiazulene | -6.765 | -3.375 | -5.065 | GN1077 | N2 |
| Water | -3.296 | 0.172 | -1.586 | GN1077 | N2 |
| Isoamyl alcohol | -16.891 | -13.635 | -15.318 | GN1077 | N2 |
| Isoquinoline | -7.372 | -3.885 | -5.597 | GN1077 | N2 |
| L-Mimosine | -5.873 | -2.464 | -4.141 | GN1077 | N2 |
| Lapachol | -6.107 | -2.442 | -4.266 | GN1077 | N2 |
| Leonurine | -6.601 | -2.893 | -4.781 | GN1077 | N2 |
| Limonin | -4.947 | -1.293 | -3.106 | GN1077 | N2 |
| Methyl palmitate | -0.995 | 2.428 | 0.735 | GN1077 | N2 |
| Oleanolic acid | -0.559 | 3.098 | 1.202 | GN1077 | N2 |
| Paeoniflorin | -8.606 | -4.582 | -6.588 | GN1077 | N2 |
| Phenylacetylene | -9.823 | -5.740 | -7.810 | GN1077 | N2 |
| Phytol | 2.564 | 6.239 | 4.456 | GN1077 | N2 |
| Piperitenone | -5.037 | -1.583 | -3.348 | GN1077 | N2 |
| Piperonyl alcohol | -5.721 | -2.376 | -4.103 | GN1077 | N2 |
| Sabinene | -4.946 | -1.221 | -3.183 | GN1077 | N2 |
| Salvinorin A propionate | 0.379 | 3.847 | 2.095 | GN1077 | N2 |
| Sinomenine hydrochloride | -5.387 | -1.905 | -3.698 | GN1077 | N2 |
| Solasodine | -4.383 | -0.653 | -2.561 | GN1077 | N2 |
| Spinosad | 2.014 | 5.509 | 3.717 | GN1077 | N2 |
| Thiophene | -12.779 | -9.263 | -11.082 | GN1077 | N2 |
| Ursolic acid | -1.001 | 2.501 | 0.708 | GN1077 | N2 |
| p-Tolualdehyde | -4.384 | -0.951 | -2.667 | GN1077 | N2 |
| alpha-Phellandrene | -8.231 | -4.663 | -6.446 | GN1077 | N2 |
| (-)-Huperzine A | -2.809 | 1.108 | -0.871 | PR678 | N2 |

| | | | | | |
|---|---|---|---|---|---|
| 1-octanol | 1.517 | 5.433 | 3.493 | PR678 | N2 |
| 2,3-Dihydrobenzofuran | -4.525 | -0.344 | -2.403 | PR678 | N2 |
| 2,5-Dihydroxybenzoic acid | -3.010 | 0.670 | -1.206 | PR678 | N2 |
| 2-Methyl-1-butanol | -12.184 | -7.739 | -9.990 | PR678 | N2 |
| 2-nonanone | -3.715 | 0.142 | -1.750 | PR678 | N2 |
| Acetophenone | -11.459 | -7.467 | -9.394 | PR678 | N2 |
| Anisole | 1.402 | 5.390 | 3.427 | PR678 | N2 |
| Camphor | 2.323 | 6.192 | 4.268 | PR678 | N2 |
| Carnosol | 2.075 | 6.238 | 4.195 | PR678 | N2 |
| Cinnamyl alcohol | -0.030 | 4.002 | 2.010 | PR678 | N2 |
| Coumaran | -2.388 | 1.468 | -0.494 | PR678 | N2 |
| Daucosterol | 4.115 | 8.281 | 6.238 | PR678 | N2 |
| Diacetyl | 2.148 | 5.876 | 4.005 | PR678 | N2 |
| Ellagic acid | 5.569 | 9.359 | 7.484 | PR678 | N2 |
| Ethyl p-methoxycinnamate | 0.070 | 4.004 | 2.059 | PR678 | N2 |
| Ethyl palmitate | -2.587 | 1.507 | -0.502 | PR678 | N2 |
| Furfural | -7.417 | -3.388 | -5.379 | PR678 | N2 |
| Guaiazulene | -4.775 | -0.936 | -2.906 | PR678 | N2 |
| Water | 1.276 | 5.399 | 3.397 | PR678 | N2 |
| Isoamyl alcohol | -12.602 | -8.954 | -10.806 | PR678 | N2 |
| Isoquinoline | -4.569 | -0.603 | -2.518 | PR678 | N2 |
| L-Mimosine | -4.180 | -0.435 | -2.327 | PR678 | N2 |
| Lapachol | -1.057 | 3.038 | 1.005 | PR678 | N2 |
| Leonurine | -3.443 | 0.623 | -1.417 | PR678 | N2 |
| Limonin | -2.736 | 1.142 | -0.779 | PR678 | N2 |
| Methyl palmitate | 3.110 | 6.904 | 5.036 | PR678 | N2 |
| Oleanolic acid | 4.736 | 9.125 | 6.850 | PR678 | N2 |
| Paeoniflorin | -4.926 | -0.601 | -2.790 | PR678 | N2 |
| Phenylacetylene | -5.255 | -0.906 | -3.118 | PR678 | N2 |
| Phytol | 8.329 | 12.420 | 10.411 | PR678 | N2 |
| Piperitenone | 1.115 | 4.972 | 3.102 | PR678 | N2 |
| Piperonyl alcohol | -1.424 | 2.442 | 0.564 | PR678 | N2 |
| Sabinene | -0.597 | 3.329 | 1.345 | PR678 | N2 |
| Salvinorin A propionate | 4.027 | 7.908 | 6.008 | PR678 | N2 |
| Sinomenine hydrochloride | -0.508 | 3.345 | 1.451 | PR678 | N2 |
| Solasodine | -1.131 | 2.879 | 0.919 | PR678 | N2 |
| Spinosad | 3.095 | 6.850 | 4.988 | PR678 | N2 |
| Thiophene | -8.857 | -4.802 | -6.835 | PR678 | N2 |
| Ursolic acid | 3.812 | 7.798 | 5.826 | PR678 | N2 |
| p-Tolualdehyde | -4.845 | -1.121 | -2.989 | PR678 | N2 |
| alpha-Phellandrene | -4.737 | -0.741 | -2.724 | PR678 | N2 |
| (-)-Huperzine A | -5.213 | -1.065 | -3.095 | CX10 | N2 |
| 1-octanol | 4.203 | 8.333 | 6.273 | CX10 | N2 |
| 2,3-Dihydrobenzofuran | -1.635 | 2.671 | 0.541 | CX10 | N2 |

| | | | | | |
|---|---|---|---|---|---|
| 2,5-Dihydroxybenzoic acid | -6.129 | -1.903 | -3.983 | CX10 | N2 |
| 2-Methyl-1-butanol | -3.197 | 1.362 | -0.928 | CX10 | N2 |
| 2-nonanone | -1.315 | 3.008 | 0.831 | CX10 | N2 |
| Acetophenone | -6.177 | -2.017 | -4.099 | CX10 | N2 |
| Anisole | -0.221 | 4.231 | 1.970 | CX10 | N2 |
| Camphor | 2.280 | 6.359 | 4.324 | CX10 | N2 |
| Carnosol | -3.022 | 1.225 | -0.856 | CX10 | N2 |
| Cinnamyl alcohol | -3.580 | 0.644 | -1.418 | CX10 | N2 |
| Coumaran | -4.180 | -0.009 | -2.130 | CX10 | N2 |
| Daucosterol | -0.144 | 4.308 | 2.080 | CX10 | N2 |
| Diacetyl | -6.107 | -1.996 | -4.021 | CX10 | N2 |
| Ellagic acid | 2.735 | 6.760 | 4.738 | CX10 | N2 |
| Ethyl p-methoxycinnamate | 0.685 | 4.852 | 2.777 | CX10 | N2 |
| Ethyl palmitate | -4.409 | -0.031 | -2.186 | CX10 | N2 |
| Furfural | -2.271 | 2.250 | -0.098 | CX10 | N2 |
| Guaiazulene | -8.349 | -4.327 | -6.332 | CX10 | N2 |
| Water | -0.388 | 3.812 | 1.653 | CX10 | N2 |
| Isoamyl alcohol | -4.732 | -0.853 | -2.780 | CX10 | N2 |
| Isoquinoline | -4.141 | 0.103 | -2.097 | CX10 | N2 |
| L-Mimosine | -5.719 | -1.428 | -3.598 | CX10 | N2 |
| Lapachol | -4.117 | 0.182 | -1.994 | CX10 | N2 |
| Leonurine | -6.955 | -2.730 | -4.797 | CX10 | N2 |
| Limonin | -2.688 | 1.552 | -0.574 | CX10 | N2 |
| Methyl palmitate | 1.487 | 5.699 | 3.565 | CX10 | N2 |
| Oleanolic acid | -2.342 | 2.020 | -0.108 | CX10 | N2 |
| Paeoniflorin | -3.276 | 1.384 | -0.883 | CX10 | N2 |
| Phenylacetylene | -3.213 | 1.545 | -0.794 | CX10 | N2 |
| Phytol | 4.498 | 9.074 | 6.754 | CX10 | N2 |
| Piperitenone | -0.086 | 3.927 | 1.956 | CX10 | N2 |
| Piperonyl alcohol | -8.682 | -4.569 | -6.562 | CX10 | N2 |
| Sabinene | -2.479 | 1.883 | -0.265 | CX10 | N2 |
| Salvinorin A propionate | 0.553 | 4.783 | 2.685 | CX10 | N2 |
| Sinomenine hydrochloride | -5.159 | -0.922 | -3.047 | CX10 | N2 |
| Solasodine | -6.434 | -2.189 | -4.364 | CX10 | N2 |
| Spinosad | 1.710 | 5.986 | 3.790 | CX10 | N2 |
| Thiophene | -10.251 | -5.991 | -8.195 | CX10 | N2 |
| Ursolic acid | 0.150 | 4.442 | 2.341 | CX10 | N2 |
| p-Tolualdehyde | -2.042 | 2.114 | -0.036 | CX10 | N2 |
| alpha-Phellandrene | -5.219 | -1.024 | -3.068 | CX10 | N2 |
| (-)-Huperzine A | -4.772 | -1.283 | -2.941 | PR678 | GN1077 |
| 1-octanol | 2.371 | 5.513 | 3.912 | PR678 | GN1077 |
| 2,3-Dihydrobenzofuran | -9.946 | -6.539 | -8.216 | PR678 | GN1077 |
| 2,5-Dihydroxybenzoic acid | -3.459 | -0.520 | -2.015 | PR678 | GN1077 |
| 2-Methyl-1-butanol | -5.712 | -1.911 | -3.813 | PR678 | GN1077 |

| | | | | | |
|---|---|---|---|---|---|
| 2-nonanone | 2.761 | 5.709 | 4.217 | PR678 | GN1077 |
| Acetophenone | -0.111 | 3.047 | 1.492 | PR678 | GN1077 |
| Anisole | -8.630 | -5.376 | -7.088 | PR678 | GN1077 |
| Camphor | -3.367 | 0.009 | -1.725 | PR678 | GN1077 |
| Carnosol | -8.770 | -5.264 | -7.094 | PR678 | GN1077 |
| Cinnamyl alcohol | -7.148 | -3.597 | -5.352 | PR678 | GN1077 |
| Coumaran | -7.063 | -3.995 | -5.543 | PR678 | GN1077 |
| Daucosterol | -4.257 | -0.676 | -2.487 | PR678 | GN1077 |
| Diacetyl | -14.111 | -11.054 | -12.577 | PR678 | GN1077 |
| Ellagic acid | -6.234 | -3.106 | -4.739 | PR678 | GN1077 |
| Ethyl p-methoxycinnamate | -1.374 | 1.684 | 0.184 | PR678 | GN1077 |
| Ethyl palmitate | -5.069 | -1.838 | -3.393 | PR678 | GN1077 |
| Furfural | -1.545 | 1.674 | 0.020 | PR678 | GN1077 |
| Guaiazulene | -3.798 | -0.470 | -2.159 | PR678 | GN1077 |
| Water | -6.680 | -3.359 | -4.983 | PR678 | GN1077 |
| Isoamyl alcohol | -6.042 | -2.955 | -4.512 | PR678 | GN1077 |
| Isoquinoline | -4.604 | -1.533 | -3.079 | PR678 | GN1077 |
| L-Mimosine | -3.361 | -0.275 | -1.814 | PR678 | GN1077 |
| Lapachol | -6.917 | -3.553 | -5.271 | PR678 | GN1077 |
| Leonurine | -5.000 | -1.681 | -3.364 | PR678 | GN1077 |
| Limonin | -3.985 | -0.714 | -2.327 | PR678 | GN1077 |
| Methyl palmitate | -5.803 | -2.732 | -4.301 | PR678 | GN1077 |
| Oleanolic acid | -7.381 | -3.858 | -5.648 | PR678 | GN1077 |
| Paeoniflorin | -5.495 | -2.068 | -3.799 | PR678 | GN1077 |
| Phenylacetylene | -6.283 | -3.075 | -4.692 | PR678 | GN1077 |
| Phytol | -7.554 | -4.319 | -5.955 | PR678 | GN1077 |
| Piperitenone | -8.009 | -4.797 | -6.449 | PR678 | GN1077 |
| Piperonyl alcohol | -6.288 | -2.974 | -4.667 | PR678 | GN1077 |
| Sabinene | -6.194 | -2.885 | -4.528 | PR678 | GN1077 |
| Salvinorin A propionate | -5.504 | -2.305 | -3.913 | PR678 | GN1077 |
| Sinomenine hydrochloride | -6.658 | -3.583 | -5.149 | PR678 | GN1077 |
| Solasodine | -5.099 | -1.824 | -3.480 | PR678 | GN1077 |
| Spinosad | -2.824 | 0.308 | -1.271 | PR678 | GN1077 |
| Thiophene | -5.878 | -2.674 | -4.247 | PR678 | GN1077 |
| Ursolic acid | -6.774 | -3.432 | -5.118 | PR678 | GN1077 |
| p-Tolualdehyde | -1.161 | 1.777 | 0.322 | PR678 | GN1077 |
| alpha-Phellandrene | -5.241 | -2.219 | -3.721 | PR678 | GN1077 |
| (-)-Huperzine A | -2.618 | 1.119 | -0.717 | CX10 | GN1077 |
| 1-octanol | -0.646 | 2.898 | 1.133 | CX10 | GN1077 |
| 2,3-Dihydrobenzofuran | -12.855 | -9.444 | -11.160 | CX10 | GN1077 |
| 2,5-Dihydroxybenzoic acid | -0.970 | 2.539 | 0.762 | CX10 | GN1077 |
| 2-Methyl-1-butanol | -14.822 | -10.784 | -12.874 | CX10 | GN1077 |
| 2-nonanone | -0.170 | 3.413 | 1.635 | CX10 | GN1077 |
| Acetophenone | -5.537 | -2.078 | -3.802 | CX10 | GN1077 |

| | | | | | |
|---|---|---|---|---|---|
| Anisole | -7.440 | -3.651 | -5.631 | CX10 | GN1077 |
| Camphor | -3.642 | -0.042 | -1.782 | CX10 | GN1077 |
| Carnosol | -3.848 | -0.309 | -2.043 | CX10 | GN1077 |
| Cinnamyl alcohol | -3.895 | 0.026 | -1.925 | CX10 | GN1077 |
| Coumaran | -5.670 | -2.195 | -3.907 | CX10 | GN1077 |
| Daucosterol | -0.218 | 3.641 | 1.671 | CX10 | GN1077 |
| Diacetyl | -6.293 | -2.774 | -4.551 | CX10 | GN1077 |
| Ellagic acid | -3.698 | -0.271 | -1.993 | CX10 | GN1077 |
| Ethyl p-methoxycinnamate | -2.211 | 1.210 | -0.534 | CX10 | GN1077 |
| Ethyl palmitate | -3.598 | 0.060 | -1.709 | CX10 | GN1077 |
| Furfural | -7.132 | -3.403 | -5.261 | CX10 | GN1077 |
| Guaiazulene | -0.554 | 3.086 | 1.267 | CX10 | GN1077 |
| Water | -4.998 | -1.528 | -3.238 | CX10 | GN1077 |
| Isoamyl alcohol | -14.353 | -10.877 | -12.539 | CX10 | GN1077 |
| Isoquinoline | -5.278 | -1.825 | -3.500 | CX10 | GN1077 |
| L-Mimosine | -2.437 | 1.233 | -0.543 | CX10 | GN1077 |
| Lapachol | -4.096 | -0.489 | -2.272 | CX10 | GN1077 |
| Leonurine | -1.738 | 1.729 | 0.016 | CX10 | GN1077 |
| Limonin | -4.352 | -0.673 | -2.533 | CX10 | GN1077 |
| Methyl palmitate | -4.686 | -1.065 | -2.830 | CX10 | GN1077 |
| Oleanolic acid | -0.549 | 3.213 | 1.310 | CX10 | GN1077 |
| Paeoniflorin | -7.549 | -3.719 | -5.705 | CX10 | GN1077 |
| Phenylacetylene | -8.885 | -5.164 | -7.016 | CX10 | GN1077 |
| Phytol | -4.060 | -0.439 | -2.298 | CX10 | GN1077 |
| Piperitenone | -7.161 | -3.527 | -5.304 | CX10 | GN1077 |
| Piperonyl alcohol | 0.671 | 4.304 | 2.460 | CX10 | GN1077 |
| Sabinene | -4.740 | -1.101 | -2.918 | CX10 | GN1077 |
| Salvinorin A propionate | -2.434 | 1.121 | -0.589 | CX10 | GN1077 |
| Sinomenine hydrochloride | -2.388 | 1.127 | -0.651 | CX10 | GN1077 |
| Solasodine | -0.057 | 3.643 | 1.802 | CX10 | GN1077 |
| Spinosad | -1.881 | 1.756 | -0.073 | CX10 | GN1077 |
| Thiophene | -4.654 | -1.154 | -2.887 | CX10 | GN1077 |
| Ursolic acid | -3.456 | 0.124 | -1.633 | CX10 | GN1077 |
| p-Tolualdehyde | -4.475 | -0.856 | -2.631 | CX10 | GN1077 |
| alpha-Phellandrene | -5.144 | -1.670 | -3.378 | CX10 | GN1077 |
| (-)-Huperzine A | 0.220 | 4.187 | 2.224 | PR678 | CX10 |
| 1-octanol | -4.800 | -0.794 | -2.780 | PR678 | CX10 |
| 2,3-Dihydrobenzofuran | -4.904 | -0.886 | -2.945 | PR678 | CX10 |
| 2,5-Dihydroxybenzoic acid | 0.804 | 4.744 | 2.777 | PR678 | CX10 |
| 2-Methyl-1-butanol | -11.208 | -7.054 | -9.061 | PR678 | CX10 |
| 2-nonanone | -4.520 | -0.652 | -2.581 | PR678 | CX10 |
| Acetophenone | -7.218 | -3.342 | -5.294 | PR678 | CX10 |
| Anisole | -0.673 | 3.477 | 1.456 | PR678 | CX10 |
| Camphor | -2.101 | 1.970 | -0.057 | PR678 | CX10 |

| | | | | | |
|---|---|---|---|---|---|
| Carnosol | 2.982 | 7.124 | 5.051 | PR678 | CX10 |
| Cinnamyl alcohol | 1.390 | 5.459 | 3.427 | PR678 | CX10 |
| Coumaran | -0.321 | 3.635 | 1.636 | PR678 | CX10 |
| Daucosterol | 2.109 | 6.191 | 4.158 | PR678 | CX10 |
| Diacetyl | 6.081 | 10.037 | 8.026 | PR678 | CX10 |
| Ellagic acid | 0.781 | 4.800 | 2.746 | PR678 | CX10 |
| Ethyl p-methoxycinnamate | -2.571 | 1.156 | -0.717 | PR678 | CX10 |
| Ethyl palmitate | -0.395 | 3.780 | 1.685 | PR678 | CX10 |
| Furfural | -7.343 | -3.180 | -5.281 | PR678 | CX10 |
| Guaiazulene | 1.403 | 5.540 | 3.426 | PR678 | CX10 |
| Water | -0.347 | 3.850 | 1.744 | PR678 | CX10 |
| Isoamyl alcohol | -9.929 | -6.063 | -8.026 | PR678 | CX10 |
| Isoquinoline | -2.388 | 1.458 | -0.420 | PR678 | CX10 |
| L-Mimosine | -0.761 | 3.221 | 1.271 | PR678 | CX10 |
| Lapachol | 1.013 | 4.985 | 2.999 | PR678 | CX10 |
| Leonurine | 1.325 | 5.447 | 3.380 | PR678 | CX10 |
| Limonin | -2.115 | 1.766 | -0.205 | PR678 | CX10 |
| Methyl palmitate | -0.473 | 3.347 | 1.471 | PR678 | CX10 |
| Oleanolic acid | 4.797 | 8.986 | 6.958 | PR678 | CX10 |
| Paeoniflorin | -3.982 | 0.135 | -1.906 | PR678 | CX10 |
| Phenylacetylene | -4.399 | -0.214 | -2.324 | PR678 | CX10 |
| Phytol | 1.591 | 5.681 | 3.657 | PR678 | CX10 |
| Piperitenone | -0.810 | 3.189 | 1.145 | PR678 | CX10 |
| Piperonyl alcohol | 5.147 | 9.211 | 7.126 | PR678 | CX10 |
| Sabinene | -0.340 | 3.541 | 1.610 | PR678 | CX10 |
| Salvinorin A propionate | 1.233 | 5.282 | 3.323 | PR678 | CX10 |
| Sinomenine hydrochloride | 2.538 | 6.469 | 4.499 | PR678 | CX10 |
| Solasodine | 3.299 | 7.206 | 5.282 | PR678 | CX10 |
| Spinosad | -0.799 | 3.237 | 1.198 | PR678 | CX10 |
| Thiophene | -0.665 | 3.438 | 1.360 | PR678 | CX10 |
| Ursolic acid | 1.355 | 5.499 | 3.485 | PR678 | CX10 |
| p-Tolualdehyde | -4.820 | -0.908 | -2.953 | PR678 | CX10 |
| alpha-Phellandrene | -1.586 | 2.268 | 0.343 | PR678 | CX10 |
